# A Comparative Analysis of Different Feature Extraction Techniques for Palm-print Images

**DOI:** 10.1101/2020.05.04.076240

**Authors:** Prateek Pratyasha, Bharati Swarnkar, Aditya Prasad Padhy

## Abstract

In this advanced decade, automatic identification of individuals is a significant achievement due to the high demand of security system. Hence, individual recognition using biometrics data is leading in the field of image processing. Although biometrics data analysis using thumb impression and finger-prints are very popular since many years, sometimes it leads to false acceptance and rejection if any physical change occurs in the finger ridges. There may be a high risk of hacking the biometrics data which is now a big challenge for cyber security employees. This paper captures the palm-print images of individuals as referred biometrics data for individual recognition. The research work is based on one of the prior issue that is feature extraction to extract the features of palm-print image such as principle lines, textures, ridges and pores etc. For this, some of the feature extraction techniques such as Derivatives of Gaussian filter (DoG), Discrete Cosine Transform (DCT), Fast Fourier Transform (FFT) and competitive coding. Two types of filters: Gaussian Filter and Gabor filter are combined with each of the feature extraction scheme for the matching of sampled image with testing image. In the result, the error rates of each of the feature extraction algorithms are compared to recognize the palm image of two different individuals.

## 1. Introduction

Biometrics is the distinctive anatomical and behavioral traits for automatic recognition of individuals. With the rapid growth of e-commerce applications, several biometric traits, including face, iris, fingerprint, palmprint, speech have been analyzed and applied to the authentication of individuals[1, 2]. The biometric use of palm prints uses ridge Patterns to identify an individual. Palms of hands epidermal ridges, thought to provide a friction surface to assist with gripping an object on surface. Palm print identification systems measure and compare ridges, lines and Minutiae found on the palm. Palm print recording and identification for law enforcement purposes has been in existence almost as long as palm prints systems are reported to comprise 30% of all crime scene marks [3]. As much as another 20% are made up of the edge of the hand, fingers between the palm and fingertips and other parts of the hand. In addition to that, with an ease of self-positioning and user-friendly nature, palmprint-based biometrics has a wide range of regular users with domestic and forensic application[4].

Some decades back, individual recognition involves only low resolution type palmprint images. For example both [5] and [6] comparatively consider the palmprint recognition algorithm by viewing the low resolution palmprint images only. Further, the Representation Based Classification (RBC) method also shows good performance in palmprint identification[7]. Additionally, the Scale Invariant Feature Transform (SIFT) which transforms image data into scale-invariant coordinates, are successfully introduced for the palmprint identification[8, 9]. However, it is observed that Lines are the basic feature of palmprint and line based methods play an important role in palmprint verification and identification. Line based methods use lines or edge detectors to extract the palmprint lines and then use them to perform palmprint verification and identification. In general, most palms have three principal lines: the heart-line, headline, and lifeline, which are the longest and widest lines in the palmprint image and have stable line shapes and positions. Thus, the principal line based method is able to provide stable performance for palmprint verification. Palmprint principal lines can be extracted by using the Gabor filter, Sobel operation, or morphological operation[10-12]. The rest of the paper is organized as: Section 2, describes the fundamental concept of palmprint images, The details of data are described in section 3. Section 4 explains various feature extraction techniques along with the filter used to extract the palmprint features. The results are analyzed in section 4 with a significant error rate detection. The conclusion of this paper is accounted in section 5.

## 2. Concept of Palmprint Images

The palmprint features have multiple levels, and different levels of features are visible in different types of palmprint images. In general, the low resolution images are of 100×100 pixels, and are texture based images, where the dark lines are more significant and visible features. Three longest and widest lines are known as principle lines and other lines are called wrinkles. Hence, the principle lines, wrinkles and texture are the most dominant features of low resolution palmprint images [13, 14]. However, the ridges of the palm cannot be visible with low resolution images, hence; there comes the requirement of high resolution images. The high resolution images are of 500×500 pixel. Generally, the ridges and ridge patterns form the fundamental features of high-resolution images [15]. Furthermore features of palmprint can be detected in a very high resolution images with 1000×1000 pixels. The palmprint images with feature levels are explained in table 1.

**Table 1.** Main Features in different resolutions of palmprint images.

| Resolution of Images | Level 1 ($100 \times 100$ pixels) | Level 2 ($500 \times 500$ pixels) | Level 3 ($1000 \times 1000$ pixels) |
| --- | --- | --- | --- |
| Features | Principle lines, texture, wrinkles | Ridges, valleys, minute points | Pores, ridge width |

It is believed that the recognition methods of palmprint images are connected with characteristics of palmprint images. Thus, it is essential to clarify all the features of types of palmprint images before extracting and matching the features. In general, palmprint images can be classified into four categories. Based on the dimension of palmprint images, they are grouped into 2-D and 3-D palmprint [16, 17]. According to the resolution, they are divided into low-resolution and high-resolution images. In addition, they can be categorized on the basis of palmprint image acquisition, and so grouped into contact-based and contactless palmprint images. Fig 5.1 shows the modes of contact-based and contactless palmprint acquisition.

**Fig 1.**
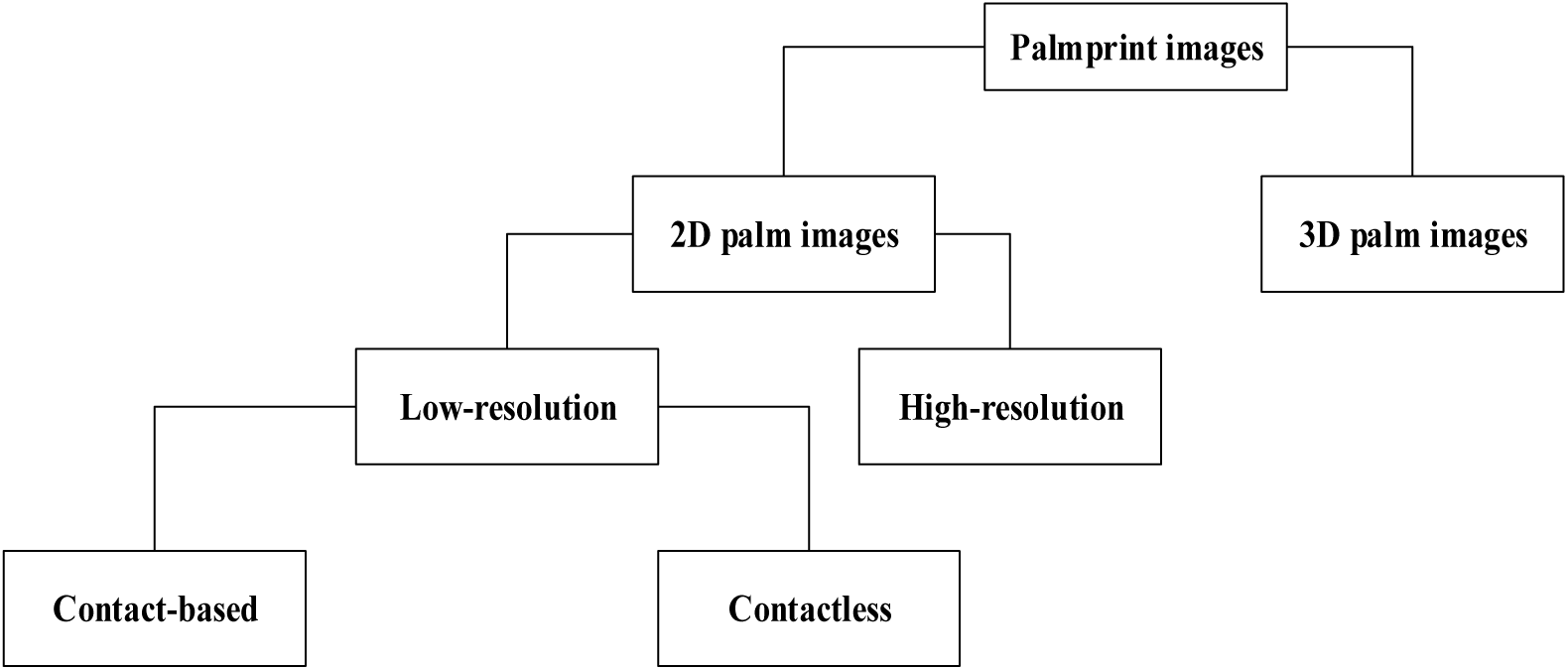
Four categories of palm images. displays the classification of various kinds of palmprint images. It is observed that the palmprint images can be better assembled in a unified framework.

## 3. Dataset description

Security and personal data is a very important topic. It covers a multitude of issues from personal, moral, to legal. Within the development of this data gathering tool security and personal privacy has been thought about carefully. This section tries to cover some of those issues. Concern is raised regarding privacy in Norway where names were asked to be deleted, leaving only Biometric data for research. Identification was permitted only by using unique identifiers. Some further checks before deployment of this generic tool will need to be made depending on the country where information is gathered and stored. In this thesis the PolyU palmprint database [“PolyU Palmprint Database,” Retrieved December 8, 2013][18] followed these rules where anonymity was guaranteed by unique identifiers that are not related to specific people.

Palmprint recognition has involves many steps to recognize a person from their palmprint images they are image reading, pre-processing, Feature Extraction, Feature matching and Decision making. In the experiments utilized the Hang Kong Polytechnic University 2D_3D_palmprint database palmprint images [19]. By taking of ROI images there is no need to perform any preprocessing. Feature Extraction method contains the methods that are utilized for extracting the features from ROI. Generally Palmprint Recognition uses five different types feature extraction algorithms, they are Subspace based approach, Texture based approach[20], Line based approach[13] and Transform based approach[21]. Subspace based approach [20] is also called as appearance based approach in the literature of palmprint recognition. Contact-based palmprint images contain visible line features. Out of them, the three principle lines such as heart line, head line and life line which are permanent for an individual. The thinner and shorter lines are sister lines or wrinkles, which are mutable. Fig 2 shows two typical contact based palmprint images.

**Fig 2.**
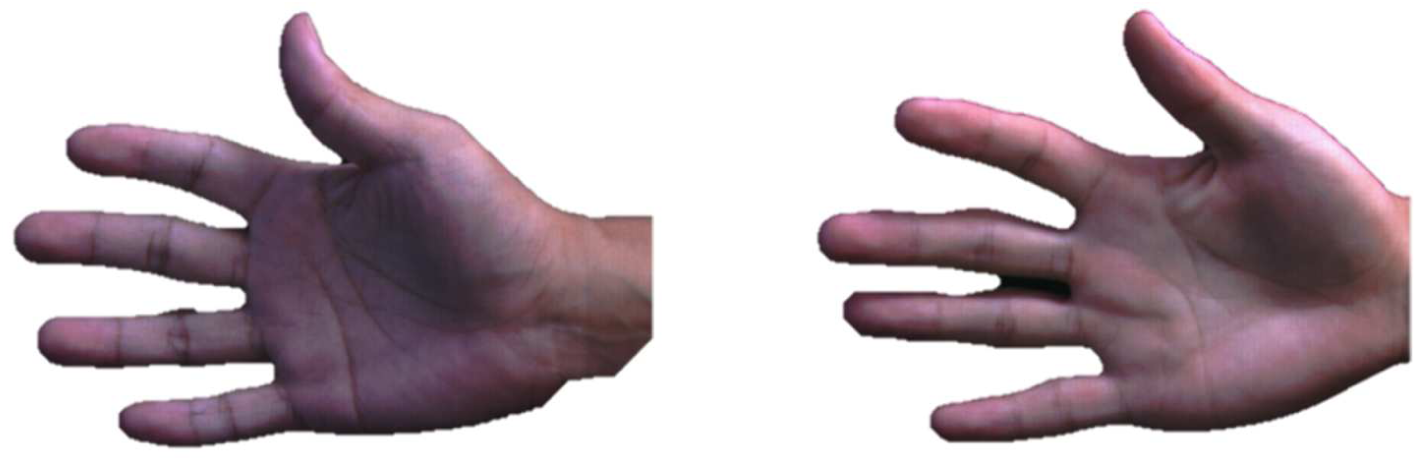
Contact-based low resolution palmprint images.

## 4. Feature Extraction Techniques of Contact-Based Palmprint Images

Feature extraction is to describe a palmprint in a wavelet feature set other than the original image. A feature with good discriminating ability should exhibit a large variance between individuals and small variance between samples from the same person. To use palmprint textures as features, transformation based approach is generally used. Among the work that appears in the literature are as follows.

### A. Difference of Gaussian Filter (DoG)

In DoG filter, a set of line detector filters based on first order and second order derivatives of a Gaussian function are present. Then, the palmprint image is convolved with these detectors. The line in Gabor filter has good performance for feature extraction however the extraction is time consuming and sensitive to nonlinear distortions and rotations. Fourier transformation includes both forward and reverse transformations of spatial to frequency and frequency to spatial conversion is possible. It is used for image enhancement, where high pass filter is used to suppose the edge lines and low pass filter used to smooth the image. But they do not provide more values to palmprint feature extraction except the frequency domain[22, 23].

### A. Discrete Cosine Transform (DCT)

Discrete Cosine Transformation (DCT) is a Fourier related transform that is equivalent to roughly twice the length of Fourier Transform but operating on real data, is used in image processing for the purpose of data compression, feature extraction. While using a large number of features for recognition it takes a large time to extract features, and it gradually decrease the recognition rate. Discrete wavelet transform for which the wavelets are discrete sampled. The wavelet transform, it is suited for analyzing images where most of the information content is represented by components. The wavelet function is designed to strike a balance between time domain and frequency domain. Wavelets are also give good feature extraction but, the selection of wavelet family and wavelet selection was difficult processor [24, 25].

### B. Fast Fourier Transform (FFT)

An image processing algorithm based on Fast Fourier transform method uses a local and global analysis for the phase unwrapping stage and gives better results than any simple Fourier Transform. It is an important image processing tool which decomposes an image into its respective sine and cosine components. The output of the transformation depicts an image in frequency domain for a spatial domain input. In frequency domain analysis, each point symbolizes a particular frequency in spatial domain image[17].

A band-pass filter of higher frequency and a cut-off filter of low frequency are set at 20 pixel and 3 pixel respectively. Thus, noise and slow variations in illumination are eliminated. This can be done by applying Inverse FFT (IFFT)[26] to the transformed image and bring it back to its reconstructed form. Fig. 5.7 depicts the Fourier transformed form of the original image and its reconstructed form. Sometimes line features are insufficient to recognize an individual with high accuracy. It may be possible that some people have same principle lines. However, the wrinkles of palmprint images usually cannot be accurately extracted for the sensitivity to noise and illumination.

### C. Competitive Coding

Competitive coding scheme [27] aims to encode the dominant orientations of palmprint lines. More concretely, let *I* (*a,b*) denote the preprocessed image, F(*a, b,θ*) is the filter with orientation*θ*, the competitive rule is defined as:

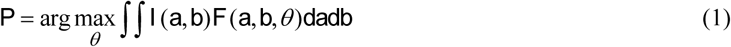

where *P* is the winning index.

According to the neuro physiological findings, the simple cells are sensitive to specific orientations with approximate bandwidths of 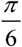 Thus, six filters with orientations 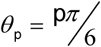, *p* = {0,1,…5} are selected for the competition.

#### Types of Filters used

##### a) Gaussian Filter

Gaussian filter is a linear time-domain filter used to blur the image. It is also filter out the noise and to reduce the contrast of the image. Mathematically, it modifies the input image by convolution by using an unique Gaussian function. The one-dimensional Gaussian function is given by the equation:

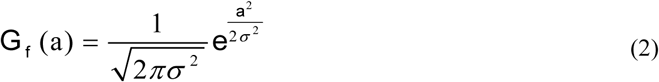

Where σ is the standard deviation and a is the sampled image.

Gaussian filter is the first step in edge detection[28]. However, it is active in smoothing the images in a better way.

In two-dimensional analysis, the overall Gaussian function is calculated by multiplying two individual Gaussian functions such as:

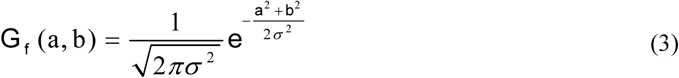

##### b) Gabor Filter

Various line and texture features are presented in palm images which carry orientation information. Since characteristics of certain cells in the visual cortex of some mammals can be approximated by Gabor filters, these filters have received considerable attention in image processing filed[29]. However, these filters can posse optimal localization properties in both spatial and frequency domain, they are well suited for texture segmentation problems. A Gabor filter can be viewed as a sinusoidal plane of particular frequency and orientation, modulated by a Gaussian envelope. It can be written as:

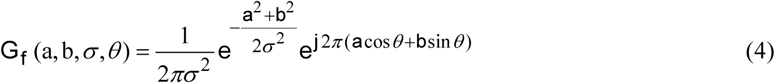

Where *θ* and σ are the orientation and bandwidth of Gabor Filter. As seen Gabor filter is complex, therefore, the output of its output also complex, and magnitude and phase of output images should be used. In this paper, we use the phase of Gabor-filtered palmprint images. We have considered wavelengths σ = {5,10} and orientations *θ* = {0,90} which results in four Gabor filters. For each ROI, we have obtained phase images.

## 5. Results and Discussion

In this paper, each of the palmprint images is matched with all of the other images in the database. The ROI and edge detection of the images are done as follows.

### FINDING ROI OF THE IMAGE

Locating the ROI of palmprint images is the most fundamental concept in biometrics and image processing. This is the primary step in developing a biometric system based on palmprint recognition. The method employed is simple and aimed to provide an efficient calculation. However, further optimizations should be possible since these requirements were not looked into in this version. The image is first smoothed by using a Gaussian filter and then by finding the centroid and updating it according to the crests and troughs[21, 29]. The distance to the centroid and minimum and maximum peaks is determined. Rectangular shape is considered for the detection of Region of Interest. Finally the sub-image of 128×128 pixels, at a certain area is cropped and resized. The original size of the images is 128 × 128 pixels. Then we cropped the main image into number of sub blocks of non-overlapping for purpose of extracting the features then the proposed approach can extract the more features using of standard deviation. The proposed algorithm for feature extraction method is

Step1: Read ROI images

Step2: Crop Original ROI image into 64×64, 32×32 and 16×16 non-overlapping and intersect of previous sub images.

Step3: Calculate, Standard Deviation for all sub images and store the values into Feature Vector.

**Fig 3.**
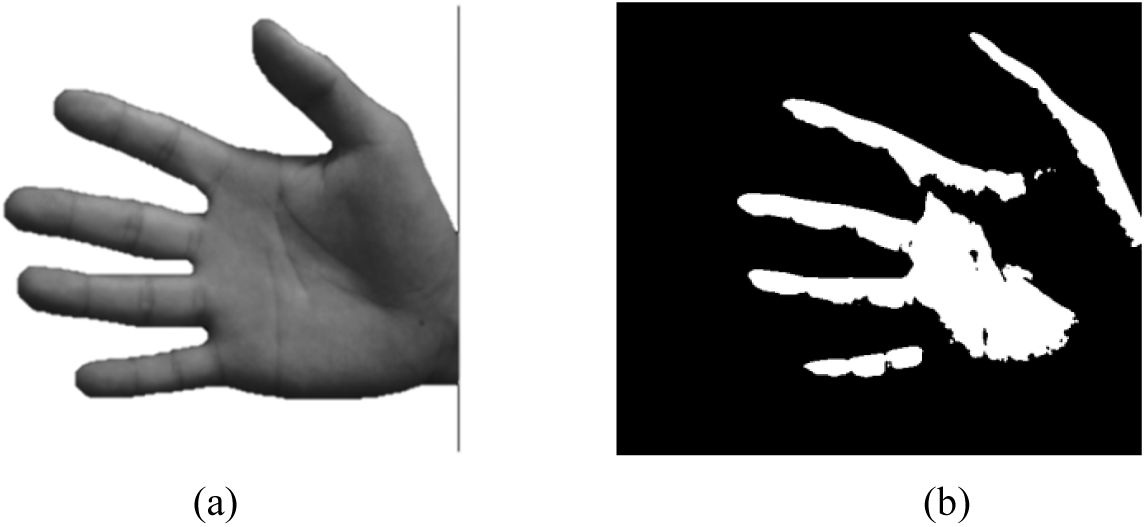
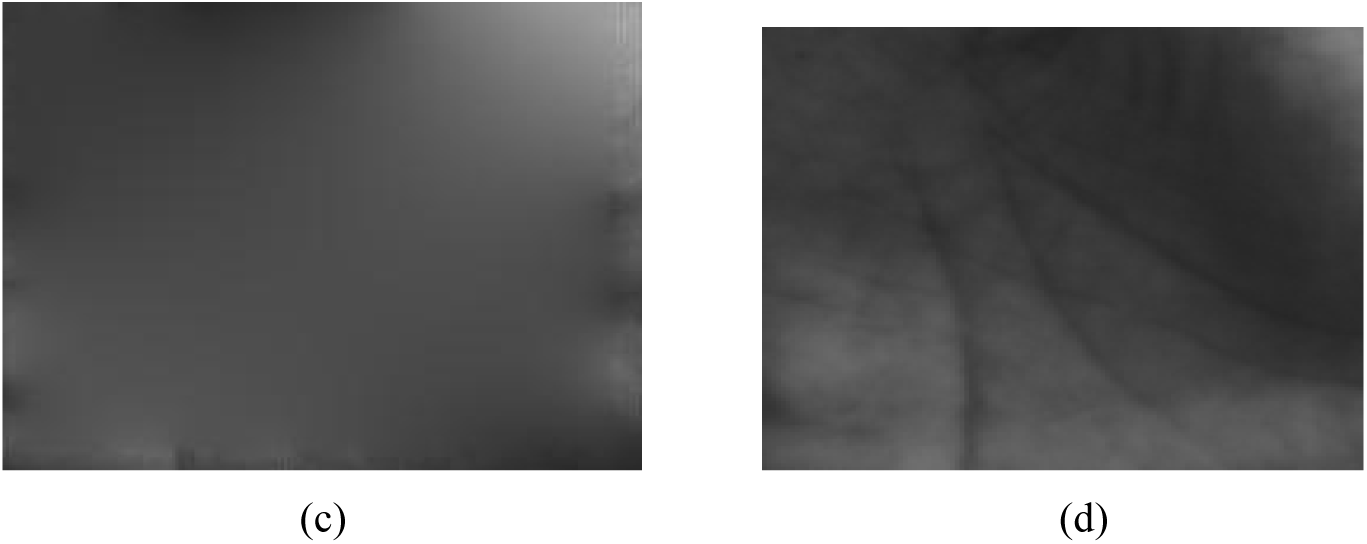
(a) Input palm image (b) Binary image (c) Boundary of image (d) Cropped ROI of palmprint image.

### SOBEL EDGE DETECTION

Image edge detection is a process of locating the edge of an image which is important in finding the approximate absolute gradient magnitude at each point I of an input grayscale image. The Sobel operator performs a 2-D spatial gradient measurement on images[30]. Transferring a 2D pixel array into statistically uncorrelated data set enhances the removal of redundant data; as a result, reduction of the amount of data is required to represent a digital image. The Sobel edge detector uses a pair of 3 x 3 convolution masks, one estimating gradient in the x-direction and the other estimating gradient in y–direction. The Sobel detector is incredibly sensitive to noise in pictures, it effectively highlight them as edges. It involves smoothing, enhancing, detection, localization. Fig 4 shows the edge detection and extracted line features from Sobel operation.

**Fig 4.**
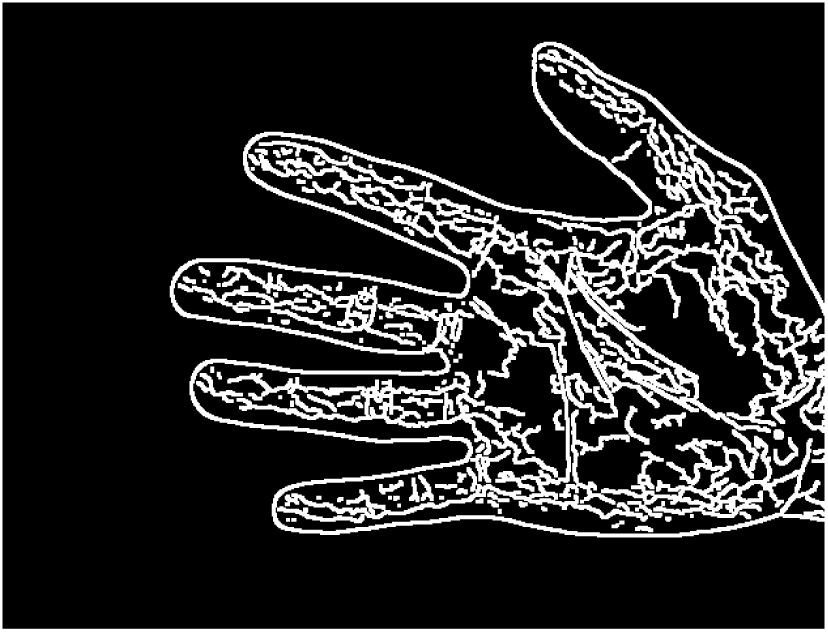
Edge detection of palmprint image.

Then different feature extraction algorithms are approached to this resized image to extract the features as well as to make the image smoother.

We have taken the 28×28 pixel of image to apply DoG method on it. Before that the coefficients of Gaussian filter was set to 0.2. To derive a smooth image, we have applied the filter coefficient to each pixel point and convolute the original image into a new image. From the new image, we have chosen the image pixels which are of approximate same weight to the original image. The, the weighted average of each new pixels of the neighborhood helped in extracting the ridges.

**Fig 5.**
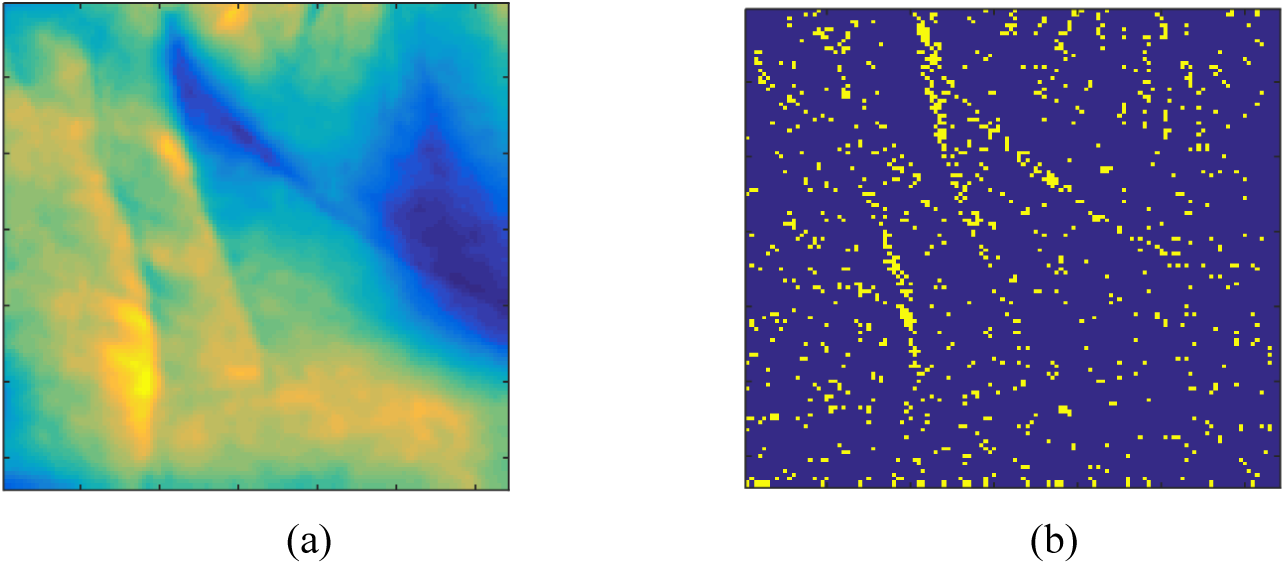
(a) Gaussian function of a palmprint image, (b) Corresponding line feature extracted from DoG filter By applying DCT to the low resolution palmprint image, the original form of the image was separated into spectral sub-bands. However, DCT is similar to Discrete Fourier Transform in converting a image from spatial domain to frequency domain. DCT is found to be more approachable than DoG because of its high degree of spectral feature extraction. And the distribution of intensity of one spectral sub-image is shown in figure 6.

**Fig 6.**
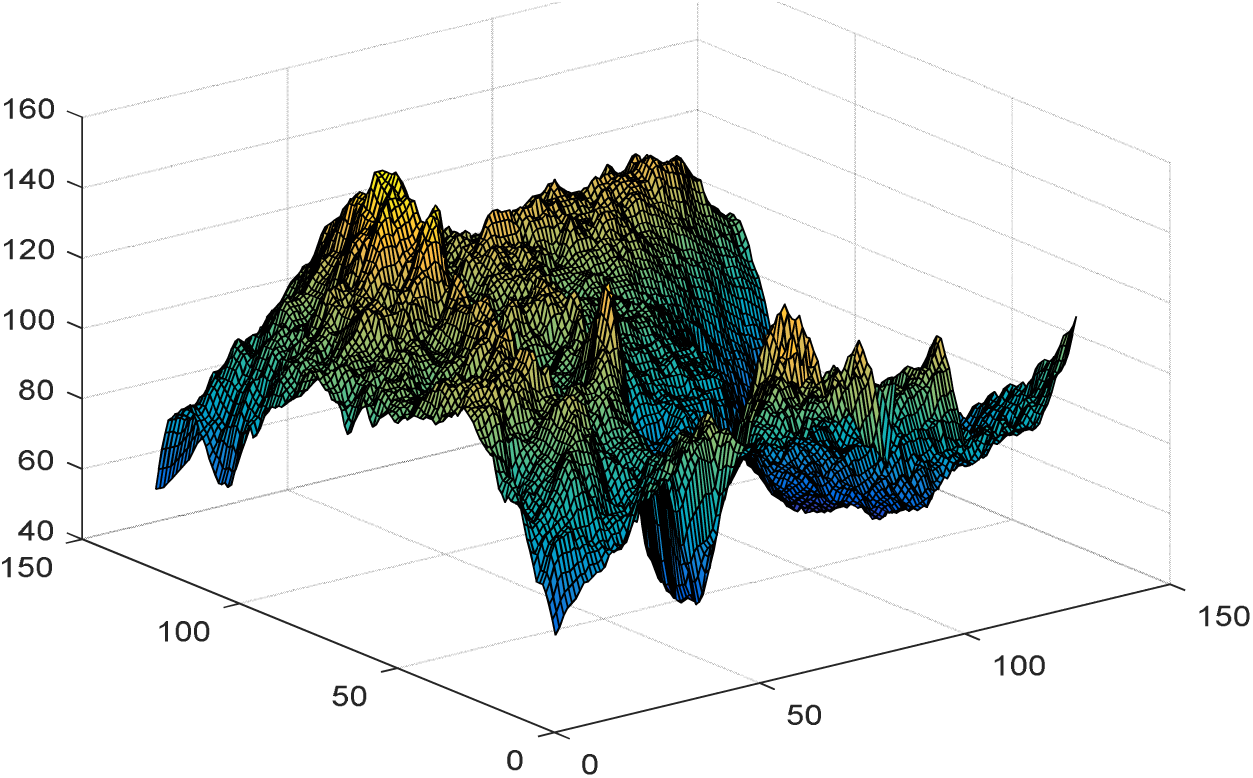
Intensity distribution of the sub-image using DCT.

FFT method is an effective reimplementation of DCT used to convert an image from spatial domain to frequency domain. The bright spot at the midpoint of the image represents the dither pattern. The frequencies corresponds to the dither pattern can be eliminated to get the encoded reconstructed image.

**Fig 7.**
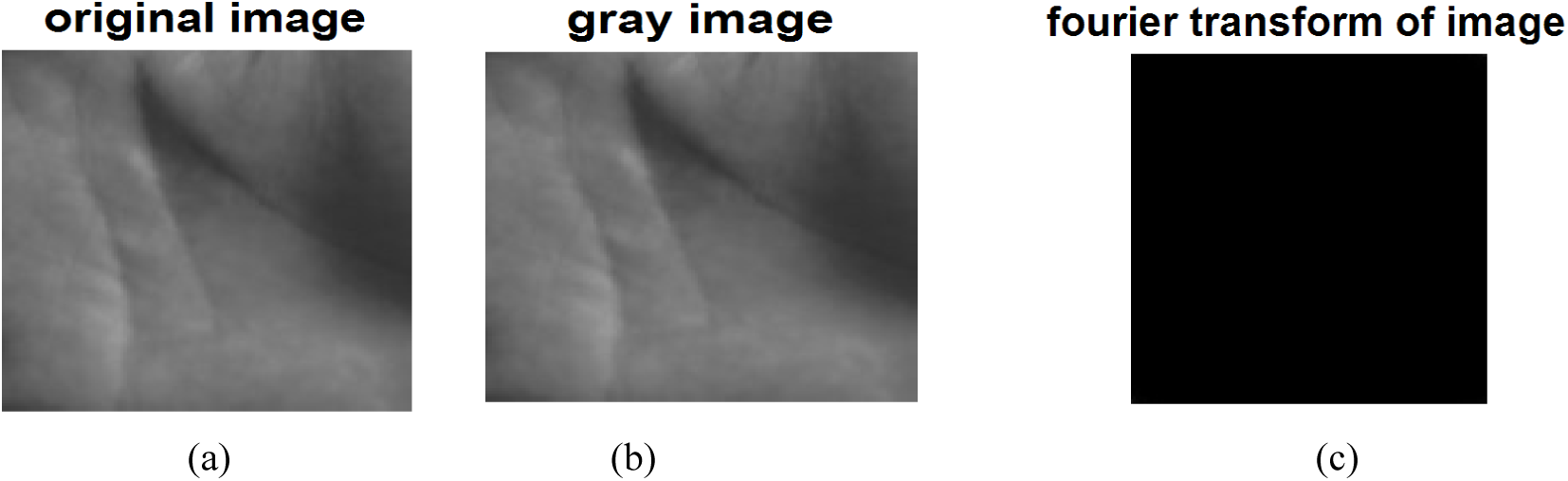
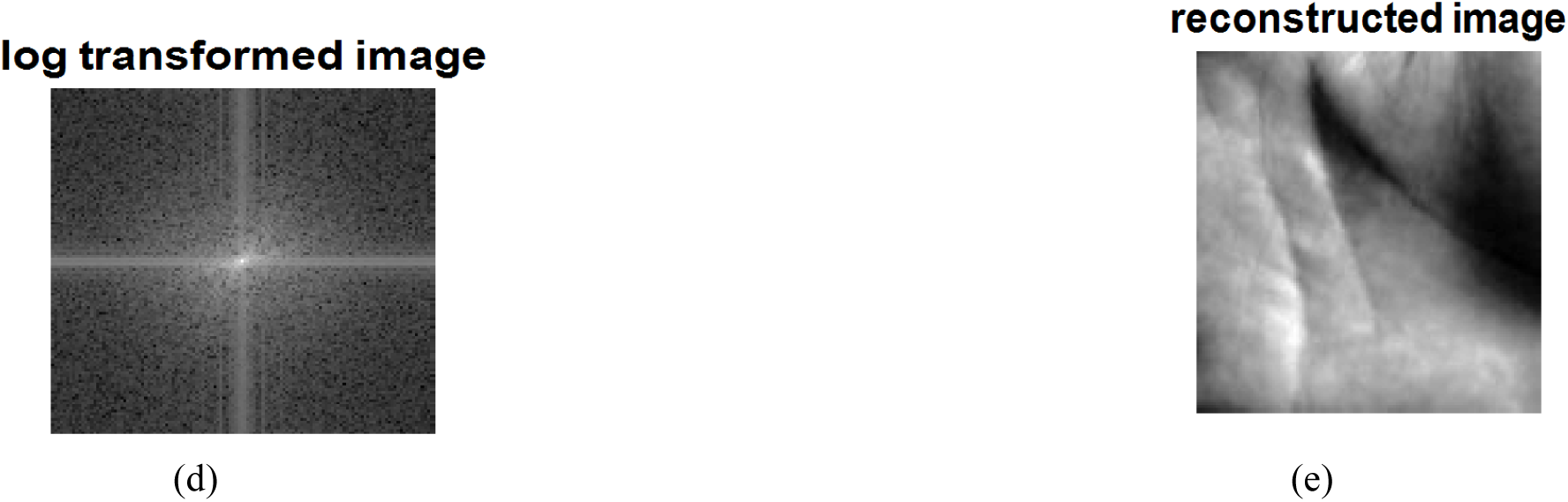
Fourier Transform of a palm image.

By using competitive coding, the two-dimensional palm image extracts the orientation related features from the images such as minute gradient lines and ridges. Because of the negative direction of palm lines, the histograms of both gradient and original images are tilted towards horizontal axis.

**Fig 8.**
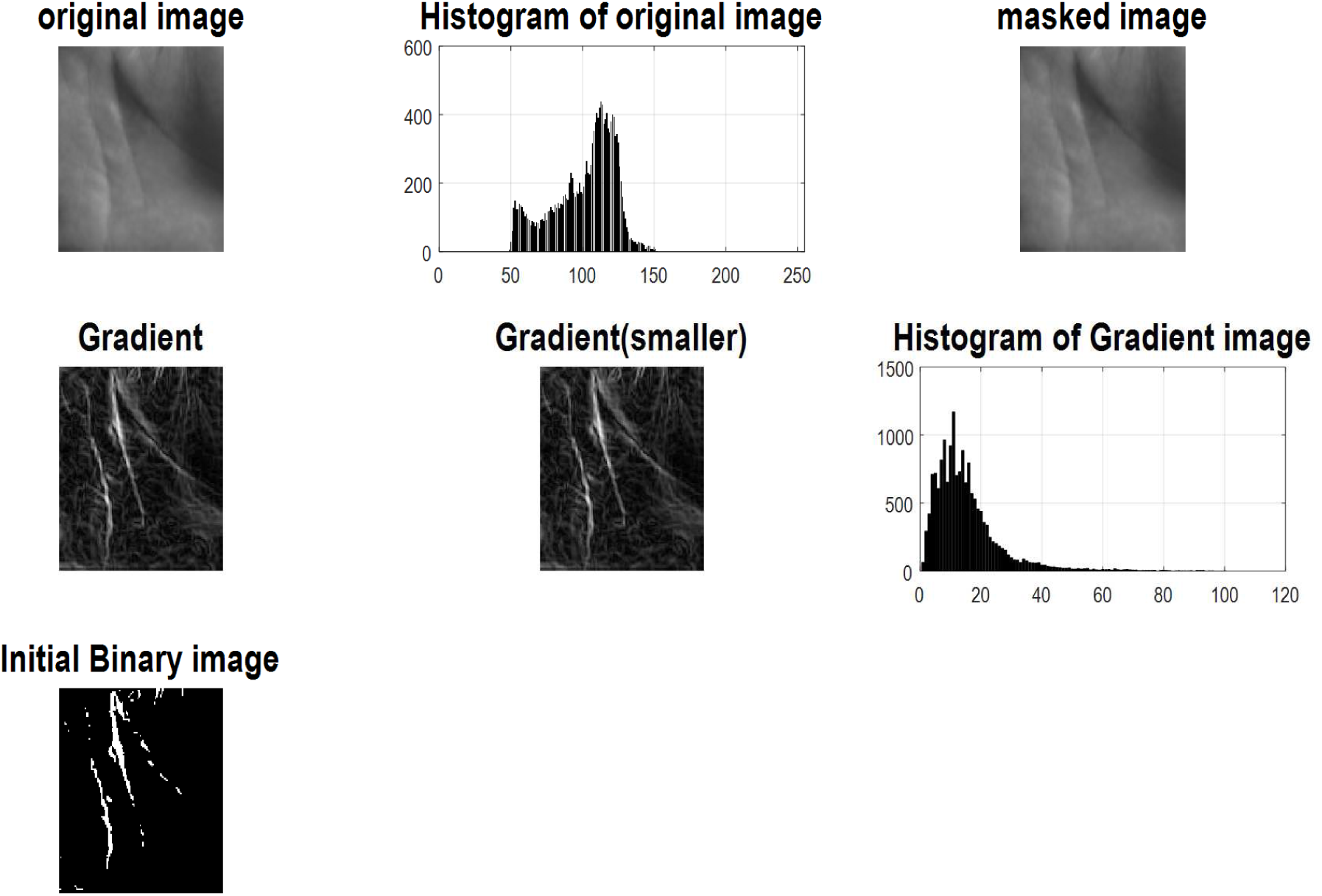
Output of gradients and its histogram by competitive code.

After finding the ROI of the original image, Gabor filter is applied to it with different orientation and wavelengths. The binary statistical image features (BSIF) of the image phase outputs are obtained. By combining different BSIF output, the average BSIF of the new palm images are computed.

**Fig 9.**
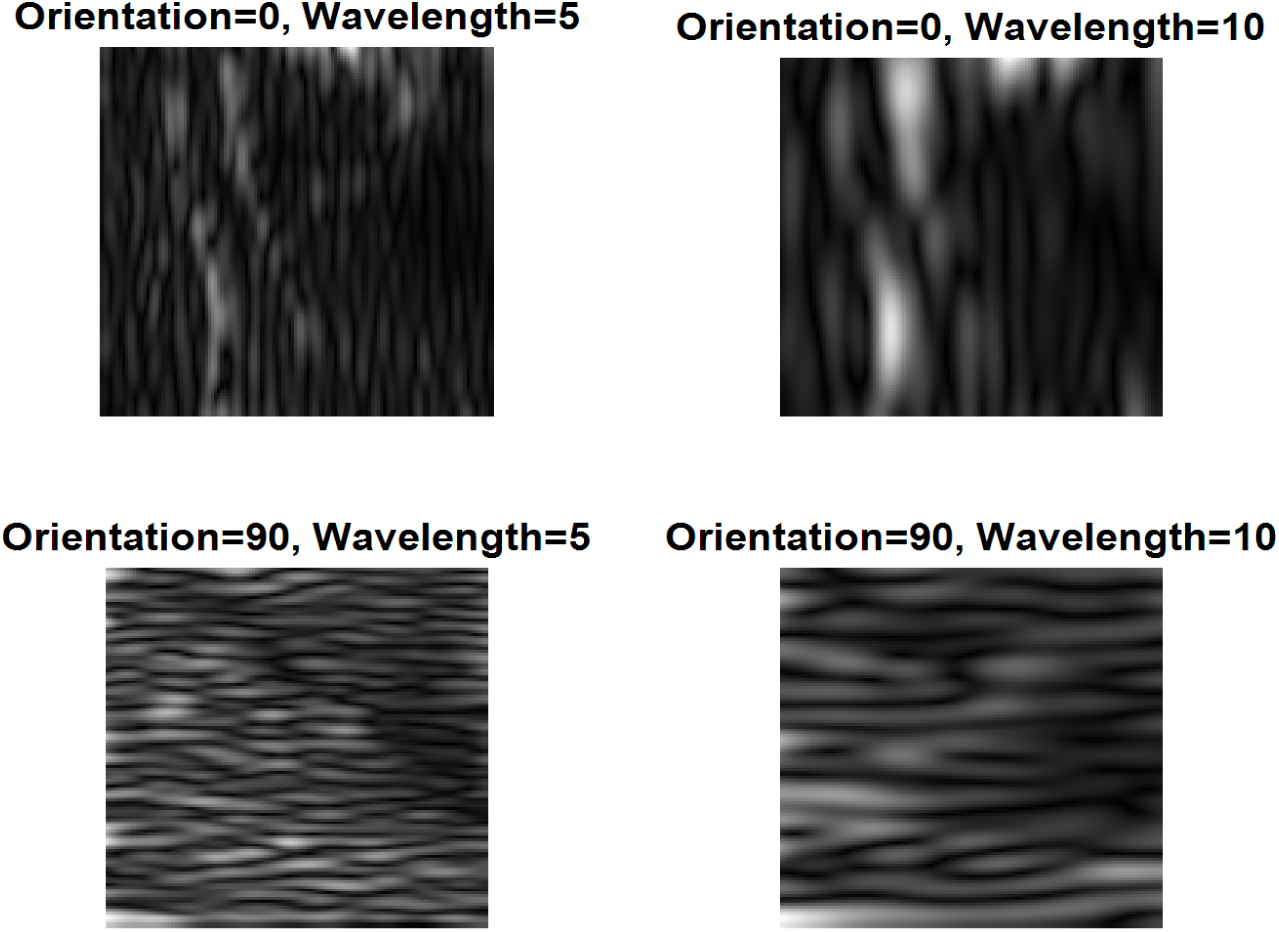
Phase output of palm image by Gabor filter.

A matching is counted as a genuine matching if two palmprint images are from the same palm; otherwise it is an imposter matching. Thus there are totally 74,068 genuine and 29,968,808 imposter matching. For all the parameters tested, the minimum Equal Error Rates (EER) of all methods are calculated. ERR is the point where False Acceptance Rate (FAR) and False Rejection Rate (FRR) overlaps which is equal to both rates[31]. For all the parameters tested, the minimum EER of all methods are listed in table 2. It is obvious that the performance of Gaussian filter is the worst of all filters.

**Table 2.** %EER obtained using different methods

| Coding Algorithms | Threshold Values | Types of filter |  |  |  |  |  |
| --- | --- | --- | --- | --- | --- | --- | --- |
|  |  | Gabor Filter |  |  | Gaussian Filter |  |  |
|  |  | FAR | FRR | EER | FAR | FRR | EER |
| **Difference of Gaussian Filter (DoG)** | 10 | 61.56 | 4 | 0.202 | 51.29 | 5.99 | 0.243 |
| **Discrete Cosine Transform (DCT)** | 15 | 55.29 | 12.95 | 0.163 | 51.96 | 27.15 | 0.219 |
| **Fast Fourier Transform (FFT)** | 20 | 38.45 | 24.75 | 0.023 | 41.09 | 38.62 | 0.186 |
| **Competitive Coding** | 25 | 5.16 | 58.2 | 0.215 | 21.87 | 48.11 | 0.257 |

From table 2, we observe that EER values obtained by Gabor filter and second derivative of Gaussian filter are comparable and the former is slightly better. Competitive coding scheme can get a smaller EER for all the three filters, which means it is superior to ordinal coding scheme. To sum up, the best result is produced by the combination of Gabor filter with competitive coding scheme. Obviously the Gaussian filter with ordinal coding scheme is the worst of all. Gaussian filter with competitive coding scheme is a little better, but still worse than other methods. Gabor filter with both coding schemes and second derivative of Gaussian filter with competitive coding scheme are comparable and among the best methods of all. Note that these results are in accordance with those of EER. The probable reason for the superiority of competitive coding scheme is that it cares only the dominant orientation of palm lines. Since most palm lines have only one dominant orientation, the response of restless dominant orientations and their relationship are not stable and prone to be disordered by noise. Thus competitive coding scheme is more robust than ordinal coding scheme when performing genuine matching, and achieves better performance accordingly.

## 6. Conclusion

We have conducted the verification experiment on PolyU Database to evaluate three filters and two coding schemes used in palmprint recognition. In summary this thesis has successfully answered the question about the role and challenges of identifying a region of interest (ROI) in palmprint recognition. This thesis also successfully quantified ROI extraction results by showing the effects on correct identification. In this paper, the new method was presented for personal identification based on palmprint. To this end, at first ROI of acquired palmprint was extracted and then was given to Gabor filter bank consists of four filters. To extract textural information from phases, we used competitive code. The competitive codes are linearly combined together with equal weights to obtain final BSIF code. Then, the normalized histogram of competitive was obtained and six features from histogram were extracted. All the coding algorithms are put individually in both Gaussian and Gabor filter to find the Equal Error Rates. The performance characteristics of ERR is independent of threshold values. After a number of evaluation, the palmprint identification rate was found to be almost 84.3%. However, there were some demerits in the entire calculation that lacks the identification rates, one of the major reason is ROI location of the sampled palm image for a better matching of image. For this we need an image of fixed pixel. Another drawback of the paper is that the computational time of most of the algorithms are more and are executed one by one. Hence, a fast computational GUI for MATLAB should be required.

